# A Targeted Mass Spectrometry Approach to detect and quantify Oxidised Phospholipids in plasma samples of Diabetic patients

**DOI:** 10.1101/741132

**Authors:** Alpesh Thakker, Corinne M Spickett, Andrew Pitt

## Abstract

Phospholipid oxidation by adventitious damage generates a wide variety of products with potentially novel biological activities that can modulate inflammatory processes associated with various diseases such as atherosclerosis, acute Pancreatitis and Type 2 diabetes. To understand the biological importance of oxidised phospholipids (OxPL) and their potential role as disease biomarkers requires precise information about the abundance of these compounds in cells and tissues. There are many chemiluminescence and spectroscopic assays available for detecting oxidised phospholipids, but they all have some limitations. Mass spectrometry coupled with liquid chromatography is a powerful and sensitive approach but its application to complex biological samples remains challenging.

The aim of this work was to develop improved methods for detection of OxPLs, specifically by using targeted mass spectrometry approaches (precursor ion [PIS] and neutral loss [NL] scanning), high resolution mass spectrometry and alternative chromatographic approaches. Initial experiments were carried out using oxidation products generated in vitro from a commercially available phosphatidylcholine (PC) and phosphatidylethanolamine (PE) mixture in order to optimise the chromatography separation parameters and mass spectrometry parameters. The chromatographic separation of oxidised phosphatidylcholines (OxPCs) and oxidised phosphatidylethanolamines (OXPEs) was evaluated using C8, C18 and C30 reverse phase, polystyrene – divinylbenzene based monolithic and mixed – mode hydrophilic interaction (HILIC) columns, interfaced with mass spectrometry. Our results suggest that the divinylbenzene based reverse phase monolithic column gave best separation of short chain OxPCs and OxPEs from long chain oxidised and native PCs and PEs.

Targeted mass spectrometric approaches for the selective identification of short chain OxPCs using PIS for m/z 184 Da and NL for m/z 34 Da for identification of hydroperoxides were tested on OxPC mixture, it enabled identification of low abundant oxidation products such as: γ-hydroxy alkenals and alkenoates and saturated aldehydes collectively termed as “short - chain oxidation products” such as PONPC, POVPC and HOOA-PC. The combination of these chromatographic and MS methods allowed identification of several oxidised molecular species in plasma of diabetic patients. Quantitative differences in oxidised products were observed in diabetic samples and the trend showed high abundance of oxidised phosphatidylcholine species in diabetic samples, compared to healthy plasma samples. However, the difference in abundance was statistically not significant when the samples were analysed using Progenesis QI software, performing global normalisation and ANOVA analysis because of inherent biological variability observed for OxPC species in samples.

## Introduction

Many diseases such as type 2 diabetes, atherosclerosis and neurodegenerative diseases including Alzheimer’s and Parkinson’s disease are associated with the inflammatory processes that is characterised by infiltration of immune cells such as neutrophils and monocytes to the site of inflammation(Fuchs, 2014). These immune cells release several enzymes like myeloperoxidase and NADPH oxidase that produce hypochlorous acid (HOCl) and superoxide radical (O_2_.-) respectively, and they undergo several reactions with other molecules and elements, forming Reactive Oxygen Species (ROS) such as hydroxyl radical (OH.). Phosphatidylcholine (PC) and phosphatidylethanolamine (PE) are the most abundant phospholipid classes present in mammalian cells, and their oxidation products have been widely studied (Birukova et al., 2013, O’Donnell, 2011). Destruction of membrane lipids involving lipid peroxidation caused by reactive oxygen species and radicals leads to compromising the integrity of its physiological role and has been correlated to many diseases and normal ageing (Aldrovandi and O’Donnell, 2013, Bochkov et al., 2010) (Reis and Spickett, 2012, Negre-Salvayre et al., 2010, Greig et al., 2012).

Moreover, a single species belonging to the glycerophospholipid group having unsaturated fatty acid esterified to the glycerol backbone can give rise to > 50 different modified species that can be grouped or classified into long chain oxidation products (where the mass of the modified compound is greater than the parent mass depending on the number of oxygen atoms attached to the species) and short chain oxidation products (formed after oxidative cleavage of long chain oxidation products)(Reis et al., 2004) (Lee et al., 2013). In addition, they can exist as many different derivatives and isomers in vivo. This can therefore, lead to formation of hundreds of different modified species from the lipidome of mammalian membrane, adding to the complexity and difficulty in characterisation and quantification of these species in biological samples (Nakanishi et al., 2009).

To understand the biological importance of oxidised phospholipids (OxPL) along with their role as a disease biomarker, information on the precise concentration of all oxidised species in biological samples must be obtained (Gruber et al., 2012). Moreover, it is important to measure simultaneously all long chain oxidation products including hydroperoxides and hydroxides, and short chain oxidation products including saturated and unsaturated aldehydes and di-carboxylic acids derivatives. This is required to investigate the dynamic assessment of oxidised phospholipids in disease state, and to examine their relevance to *in vivo* function and disease (Uchikata et al., 2012). A variety of techniques such as biological assays like Fox2 assay, Isoluminol assay has been developed to measure free radical products in disease states. Although, these methods provide useful initial insights about potential oxidative state, they suffer from inherent problems related to sensitivity and specificity of detecting individual molecular species, when applied to *in vivo* situations (Tyurina et al., 2009) (Spickett et al., 2011).

On the other hand mass spectrometry coupled with liquid chromatography as a technique to measure oxidised phospholipids has gain popularity because it measures mass –to charge ratio (*m/z*) of compounds and can distinguish or separate different molecular species based on their masses and functional chemistry, therefore, can selectively identify several different species in complex mixture simultaneously (Sandra et al., 2010, Gruber et al., 2012). Unsurprisingly, several liquid chromatography - mass spectrometry based methods have been developed for analysis of oxidised phospholipids. Most of the current methods published so far, have been either confined to small number of molecular species based on Multiple Reaction monitoring (MRM) based approach or have used high resolution mass spectrometry coupled with liquid chromatography to detect specified oxidised phospholipid species, solely based on accurate masses and/or tandem mass spectrometry (MS-MS) to obtain structural information (Adachi et al., 2004, Spickett et al., 2011). The MRM and high resolution based methods are highly sensitive and focused methods and are ideal for observing and quantifying certain pre-determined species. Nevertheless, further improved methods are still required, that is capable of quantifying multiple oxidised species in biological samples belonging to different phospholipid class. However, it is analytically impractical to measure oxidised species of all classes in a single run because of variation in charge of polar head group and extreme diversity of oxidised phospholipid species. Various reports describe the combination of either reverse phased and or normal-phase LC coupled with mass spectrometry in multi-dimensional set ups to detect and measure long chain oxidation products (Hui et al., 2010, Reis et al., 2013, Uchikata et al., 2012, Tyurina et al., 2009, Morgan et al., 2010). Our study focus on the detection of oxidised phosphatidylcholines (OxPCs), which represent an abundant class demonstrating variety of biological activities but insufficiently characterised in several inflammatory diseases. Targeted based approaches like precursor ion scanning (PIS) and neutral loss scanning (NL) developed based on the knowledge of common fragmentation patterns of a small group of molecular species, bearing a common structural motif, have been reported earlier but have not been extensively used to measure OxPC species in biological samples (Reis et al., 2013) (Spickett et al., 2011).

This article reports two different improved methods developed on a hybrid Q-Trap mass spectrometer to detect hydroperoxides and chlorohydrins respectively, using targeted approaches based on neutral loss scanning (NL) reported in (Spickett et al., 2011). Furthermore, the chromatographic separation of these diverse ranges of oxidation products formed was evaluated using several reverse phase columns and mixed mode columns. We have also attempted to assess the method for qualitative and quantitative use by running technical replicates, calculating their peak area and subsequently coefficient of variance (CV%).To validate our findings, same samples were analysed on a high resolution mass spectrometer (ABSciex 5600 Q-TOF) and data analysis was performed using narrow mass window and accurate masses with error less than 5ppm, using Progenesis QI software that is capable of performing global normalisation, baseline correction and peak picking; identification using in-house database and statistical analysis. Type 2 diabetes is a complex polygenic disorder of intermediary metabolism with pathogenesis related to insulin resistance. It has also been associated with the oxidative stress and the metabolic disorder of phospholipids caused by overproduction of ROS by mitochondria (Giacco and Brownlee, 2010). While research has been published to analyse age related glycation products associated with phospholipid and proteins, an extensive characterisation of OxPC species has not been done (Miyazawa et al., 2012, Shoji et al., 2010). Therefore, we investigated the abundance of several OxPC species in lipid extracts of plasma of diabetic and healthy patients.

## Materials and Methods

L-α phosphocholine mixture (PC mixture from egg yolk) and 2-2’-Azobis (2-methylproprionamidine).2HCl (AAPH), a water soluble radical initiator, was obtained from Sigma-Aldrich, Dorset, UK. 1, 2-ditridecanoyl-sn-glycero-3-phosphocholine (PC (13:0, 13:0)) used as internal standard and was procured from Avanti Polar Lipids, USA. All solvents (methanol, chloroform, water) were of HPLC grade and obtained from Thermo Fisher Scientific, UK. Pro-swift RP-4H (1×250mm) monolithic column was purchased from Thermo Scientific, UK for LC-MS analysis. The Luna C8 column (150mm × 1 mm), Luna C-18 column (150 × 1 mm) and C-30 column (150 × 2.1 mm) were purchased from Phenomenex, UK. Mix – mode HILIC column (300 × 2.1 mm) was procured from HiChrom, UK.

### Preparation of Vesicles

PC mixture was pre-aliqoted as 100 µg dried weight and reconstituted in 10 µl milli-Q water to make a 10mg/ml suspension. The suspension was vortexed for 1 minute followed by sonication for 30 min. The mixture was further vortexed for 1 minute to form multilamellar lipid vesicles.

### Oxidative Treatments

#### Generation of peroxidation products from PC mixture

Lipid vesicles were prepared as described above followed by addition of 100 µl of 10 mM AAPH solution. The reaction mixture was incubated for 24 hr at 37°C. The reaction was stopped by placing the vial on ice and adding 100 µl of ice cold Methanol containing 0.005% BHT. 100 ng of PC (13:0, 13:0) was added to the mixture as internal standard and lipid extraction was carried out using modified Folch method. The organic layer collected was dried under gentle stream of nitrogen and stored in −20 °C freezer until further analysis.

#### Generation of chlorohydrins products from PC mixture

The PC mixture vesicles (10mg/ml) were treated with 150 mM of sodium hypochlorite (NaOCl) at pH 6.0 for 30 min to achieve maximum conversion of native lipid to chlorohydrin. Excess hypochlorite was removed by passing the mixture through a reverse phase C18 Sep-pak cartridge (Waters, UK) and washing with an excess of water. The column was pre-conditioned with methanol (1 ml) and equilibrated with water (2 ml) prior to loading the sample and lipids were extracted with 100% methanol (0.5 ml) and 1:1 methanol/chloroform (1.5 ml). The organic solvent was dried under nitrogen and PC mixture chlorohydrins (PC Mix-ClOH) were stored in −20° C freezer until further analysis on a mass spectrometer.

### Lipid extraction after treatment

After the oxidative treatment, the oxidised PC mixture (OxPC) was supplemented with 100 ng of (13:0/13:0) PC as internal standard by adding 100 µl of 1 µg/ml (13:0/13:0) PC in methanol with BHT, followed by adding 100 µl of methanol containing 0.005% BHT. The mixture was sonicated for 15 minutes followed by adding 400 µl of chloroform and 150 µl LC-MS grade water. The mixture was vortexed and sonicated for 15 minutes followed by centrifugation at 14500g for 2 minutes. The top aqueous layer was collected and re-extracted with 400 µl of chloroform as above. The upper phase was discarded and both organic phases were combined, dried under nitrogen stream and stored at −70°C.

### Optimisation of chromatographic separation using several reverse phase columns and eluent systems

Conventional reverse phase columns like C-8 Luna column (150 × 1mm), C-18 Luna column (150 × 1mm), C-30 Luna column (150 × 2.1mm) and Proswift –RP 4H polystyrene – divinyl benzene coated monolith column (1 × 250mm); and mix mode Hichrom amino based HILIC column (150 × 3.1mm) were evaluated using combination of different eluent systems, while maintaining the linear flow rate and using mass spectrometry for detection. Different flow rates were tested for monolith column to improve chromatographic resolution of co-eluting isomeric and isobaric species.

### LC-MS analysis of peroxidation and chlorohydrins products of PC mixture

PC mixture oxidation products were separated by reverse phase chromatography on an HLPC system (Dionex ultimate 3000 system) controlled by Chromoleon software, fitted with the Proswift RP-4H column (1mm × 250mm) kept at room temperature in the column compartment. The eluent used was (A) water with 0.1% formic acid and 5mM ammonium formate and (B) methanol with 0.1% formic acid and 5mM ammonium formate. The run time for the entire LC-MS was 50 minutes and the chromatographic elution was programmed using dionex chromatographic management system where the mobile phase at 0 – 4 time was fixed at isocratic hold of 70% B followed by 3 step gradient increase to 80% B at 8 minutes, 90% B at 15 and 100% B at 20 minutes. The gradient remained constant until 38 minutes and decreased back to 70%B until 50 minutes. The flow of the mobile phase was set to 50 µl/minute. The mass spectrometry detection using targeted scanning routines was performed on a Absciex 5500 QTrap instrument and the validation of data analysis was performed on a high resolution Absciex Q-TOF instrument using accurate masses.

Eluting Oxidised PC mixture (OxPC) was detected on a Q-Trap Absciex 5500 mass spectrometer controlled with Analyst software. The targeted approaches for detection of OxPC used were Neutral loss scanning (NL) of 34 Da for mass range of 700 Da – 1000 Da and precursor ion scanning (PIS) of 184 Da for mass range 400 Da −1000 Da. The declustering potential was set to 50 V for all scans; collision energy for NL scan set to 35 eV and PIS scan set to 45 eV. Information dependent data acquisition (IDA) was used to collect MS/MS data based on following criteria: 1 most intense ion with +1 charge and minimum intensity of 1000 cps was chosen for analysis, using dynamic exclusion for 20 seconds after 2 occurrences and a fixed collision energy setting of 47 eV. Other source parameters were adjusted to give optimal response from the direct infusion of a dilute solution of standards.

For analysis of OxPC samples on high resolution Q-TOF mass spectrometry, similar chromatographic elution program was used and the Q-TOF MS survey scan was collected in positive mode from 400 Da – 1200 Da using high resolution and accumulation time of 250 ms. IDA was used to collect MS/MS data for 4 most intense ions with intensity greater than 200 cps, fixed collision energy of 45 eV and dynamic exclusion activated after 2 occurrences for 20 seconds. The experiment was repeated 3 times with each experiment performed in triplicates.

### Lipid extraction from plasma samples of diabetic patients

The plasma samples of diabetic patients (N=8) and healthy volunteers (N=8) were collected after ethics committee approval. Plasma sample (40 µl) was used for lipid extraction using the method described above using 100 ng of (13:0/13:0) PC as internal standard. The dried lipid extract was reconstituted in 1:1 methanol:chloroform mixture before preparing 10-fold dilution with starting solvent (70% Methanol with 5mM ammonium formate and 0.1 % formic acid) and analysis was performed on Qtrap 5500 followed by Q-TOF MS.

### Data Analysis and Statistical evaluation

Data analysis was performed manually using Peakview 2.0 software by calculating peak area through generating Extracted ion chromatograms (XIC) for individual masses of different oxidised species. The peak area was normalised using the peak area of the internal standard and the mean value of three technical replicates was calculated and used for relative quantification. Alternatively, the data validation and statistical evaluation was performed by automated analysis and global normalisation using Progenesis QI software.

## Results

The chromatographic separation of OxPC species was evaluated using several columns like C-8 Luna column (150 × 1mm), C-18 Luna column (150 × 1mm), C-30 Luna column (150 × 2.1mm) and Proswift –RP 4H polystyrene – divinyl benzene coated monolith column (1 × 250mm); and mix mode Hichrom amino based HILIC column (150 × 3.1mm) using different solvent systems. Extracted ion chromatogram of two representative species of short chain oxidation products, long chain oxidation products and native PCs separated on different columns and solvent system was obtained and the average time point of the elution for each group was used to evaluate the separation. The result illustrates that the polystyrene-divinylbenzene coated monolith column gave best separation of short chain oxidation products from long chain oxidation products and unmodified PCs and PEs, compared to other tested columns and solvent systems (Figure 1) (Supplementary figure 1). Furthermore, different flow rates were tested to achieve maximum chromatographic resolution for structurally similar oxidation products that co-eluted on monolithic column, and contributing to peak broadening. We found that the linear flow rate of 50 µl/minute was optimum to get best separation and sensitivity (Supplementary table 1). For all the subsequent experiments, the monolith column was used for chromatographic separation.

Non-enzymatic oxidation of PC mixture using 10 mM AAPH and incubation for 24 hrs formed variety of oxidised species. The chromatographic separation using monolith column, and detection on Q-trap mass spectrometer using targeted approaches like PIS for 184 Da for all oxidised and native species of phosphatidylcholine class, and NL 34 scan for detection of hydroperoxides in conjunction with IDA based MS/MS scans for structural information, enabled identification of more than 100 oxidised species (figure 2). The figure 2 shows the overlaid total ion chromatogram (TIC) of control and oxidised sample for PIS 184 scan and NL 34 scan. It illustrates the formation and up regulation of short chain oxidation products that eluted between 5-14 minutes, and long chain oxidation products eluting between 18-22 minutes, in oxidised sample, compared to control sample. The figure 2(c-d) illustrates the unequivocal identification of hydroperoxide species that can be achieved by NL 34 scanning. Multiple peaks may represent isobaric compounds with different chemical composition or may correspond to multiple positional isomers of the same OxPC, which elutes as clusters of poorly separated peaks. As an example illustrated in figure 2, the native di-unsaturated phosphatidylcholine: 1-plamitoyl, 2-linoleoyl-phosphatidylcholine (PLPC, m/z 758.5) after oxidation forms a di-hydroxide species, m/z 790.5 (by addition of 2(OH)), and a monohydroperoxide species, m/z 790.5 (by addition of OOH). The native phospahtidylcholine species (di-stearoyl-phosphatidylcholine, m/z 790.5) is also isobaric to the above mentioned oxidation species and these all species were observed in the PIS 184 scan. While the XIC for m/z 790.5 from PIS 184 scan showed 3 chromatographic peaks at 17.5 minutes, 19.5 minutes and 25.5 minutes, the XIC for m/z 790.5 showed a single peak at 19.5 minutes. Thus, NL 34 scans can be used to identify all hydroperoxide species.

Non-enzymatically oxidised PC mixture was analysed by same LC-MS/MS procedure to estimate the retention time of different species formed via free radical mechanism. As expected, due to oxidative fragmentation of oxidised fatty acid residue or addition of oxygen, the OxPC species eluted from the reverse-phased monolithic column significantly earlier as compared to the native phospholipids. We were able to detect more than 100 different oxidised species using our method based on targeted approaches, which were up regulated in the oxidised sample, compared to control sample (figure 3). The XIC’s were generated for the nominal masses of all previously identified OxPC species and peak area was measured. The experiment was repeated three times, measured peak area was averaged and normalised to the peak area of the internal standard and heat map was generated for mean peak area of all species in control and oxidised sample, representing colour coded abundance.

Based on previously identified and functionally characterised types of oxidation products, we measured exact masses, molecular formula, and retention time using current chromatographic condition for monolith column and detection on high resolution Q-TOF mass spectrometer was performed to validate our identification (Supplementary table 2). The list consisted of 5 different categories of oxidised PCs consisting of lyso-products, short chain oxidation products encompassing fragmented ω-terminal, fragmented α-β unsaturated aldehydes and dicarboxylic acids, non-fragmented prostane ring species, long chain oxidation products (hydroperoxides,hydroxides) and native PCs. Subsequently, the method that was developed on Q-Trap mass spectrometer, was employed to characterise the OxPC species in plasma of diabetic and healthy patients (N= 16). We were able to observe a plethora of long chain and short chain oxidation products in plasma of diabetic and healthy patients. Manual data analysis was performed using a label free approach with normalisation using the internal standard.

Although, the abundance level of some OxPC species were found to be up-regulated in diabetic samples, a statistical analysis using ANOVA showed this to be non-significant on account of inherent biological variability for these species that were found in plasma samples (figure 4). To validate our data analysis, the samples were analysed on a high resolution Q-TOF mass spectrometer and separation was achieved using similar chromatographic conditions using monolith column. The data acquired on Q-TOF MS were analysed using Progenesis QI software, which can automate the analysis and pre-processing steps and has capability to perform global normalisation for all ions. Such pre-processing steps and normalisation may eliminate any non-biological variance caused by matrix effects. The data analysis using progenesis software showed similar results and the data is summarised in (table 1).

**Table 1:** Data analysis of diabetic and healthy patients generated on high resolution Q-TOF mass spectrometer, using progenesis QI software. The raw files were uploaded onto the software, and the runs were aligned, followed by peak picking and global normalisation. The database developed in house was uploaded and the identification was carried out with mass tolerance set to 5 ppm.

| Description | Retention Time | Peak Width | Anova (p) | Max Fold Change | Highest abundance observed |
| --- | --- | --- | --- | --- | --- |
| SLPC(OH)-OOH | 19.95 | 0.56 | 0.761 | 3.07 | Healthy Plasma Samples |
| 15-J2IsoP with Palmitoyl | 19.4 | 0.53 | 0.891 | 2.57 | Diabetic Plasma samples |
| SDHPC-OOH | 19.57 | 0.51 | 0.069 | 2.47 | Diabetic Plasma samples |
| SLPC-OOH | 21.01 | 0.75 | 0.668 | 1.85 | Diabetic Plasma samples |
| SAPC-OOH | 19.28 | 2.04 | 0.125 | 1.70 | Diabetic Plasma samples |
| PLPC-OOH | 19.35 | 0.87 | 0.18 | 1.68 | Diabetic Plasma samples |
| SAPC-OOH | 21.74 | 1.81 | 0.401 | 1.58 | Diabetic Plasma samples |
| SLPC-2(OH) | 19.59 | 0.92 | 0.921 | 1.55 | Diabetic Plasma samples |
| PPPC* (Palmitoleic acid) | 21.96 | 1.66 | 0.082 | 1.52 | Diabetic Plasma samples |
| PAPC-OOH | 17.82 | 3.55 | 0.176 | 1.47 | Diabetic Plasma samples |
| PDHPC-OH | 20.27 | 2.76 | 0.152 | 1.42 | Diabetic Plasma samples |
| SLPC-OH | 20.92 | 0.92 | 0.252 | 1.31 | Diabetic Plasma samples |
| SDHPC-OH | 21.79 | 1.76 | 0.162 | 1.29 | Diabetic Plasma samples |
| SDHPC | 24.64 | 2.54 | 0.222 | 1.27 | Diabetic Plasma samples |
| PLPC(OH)-OOH | 18.01 | 1.18 | 0.538 | 1.27 | Healthy Plasma Samples |
| SAPC-OH | 21.79 | 3.09 | 0.171 | 1.25 | Diabetic Plasma samples |
| 20:4 lysoPC | 5.67 | 2.93 | 0.119 | 1.24 | Diabetic Plasma samples |
| 16:0 lysoPC | 5.56 | 2.41 | 0.062 | 1.24 | Diabetic Plasma samples |
| 18:2 lysoPC | 5.56 | 1.57 | 0.18 | 1.22 | Diabetic Plasma samples |
| PLPC-OH | 19.18 | 2.2 | 0.312 | 1.21 | Diabetic Plasma samples |
| POVPC | 6.21 | 0.89 | 0.356 | 1.21 | Diabetic Plasma samples |
| POPC-OH | 21.59 | 1.57 | 0.214 | 1.20 | Diabetic Plasma samples |
| SAPC | 24.28 | 2.67 | 0.149 | 1.17 | Diabetic Plasma samples |
| PDHPC | 23.32 | 2.4 | 0.351 | 1.16 | Diabetic Plasma samples |
| PONPC | 8.06 | 0.88 | 0.883 | 1.06 | Diabetic Plasma samples |
| PLPC | 22.58 | 2.47 | 0.509 | 1.04 | Diabetic Plasma samples |
| 18:0lysoPC | 7.7 | 2.21 | 0.926 | 1.02 | Healthy Plasma Samples |
| PAPC | 22.97 | 2.48 | 0.988 | 1.00 | Diabetic Plasma samples |

## Discussion

The major objective of this work was to develop a method using targeted approaches to detect a range of oxidised PC species within one analytical run and is sensitive enough to be translated to analyse OxPC species in different biological samples.

An important aspect of our methodology was to evaluate the separation of several oxidised species using several reverse phase columns and we found that the best separation was achieved using monolith column as compared to other conventional reverse phase columns and mix mode HILIC column. While many LC-MS based method have been reported for analysis of OxPCs, limited work was done to evaluate the separation of oxidised phospholipid species and achieve chromatographic resolution (Hui et al., 2010). Most of the method reported have concentrated on faster analysis like the method reported for analysis of OxPCs by (Gruber et al., 2012) used a C-18 core shelled column and all the species reported eluted in the first 10 minutes. However, faster analysis can compromise the chromatographic resolution and sensitivity and therefore, can affect the relative quantification process. Moreover, faster analysis of complex samples can also lead to more false positive identification because overlapping of signals of co-eluting analytes and matrix effects. Therefore, better dispersion of separation is required for analysis of samples, when the objective is to identify and characterise several species including isobaric and isomeric species in a single run.

Furthermore our work aimed to develop a method using targeted scanning approaches, which has been reported earlier but lesser exploited to characterise oxidised species in biological samples(Spickett et al., 2011) (Reis et al., 2013). With the fact, that a single poly-unsaturated phospholipid species can give rise to several oxidation products that can make the identification and quantification process of all species in a single run, a difficult task. Moreover, most of the earlier work done was reported and tested on in vitro oxidised samples and concentrated on analysis of pre-determined species. While the approach of multiple reaction monitoring (MRM) is the most used approach to quantify several pre-defined species, it has the limitations of only able to quantify certain number of species in a single run. Moreover, most of the mass spectrometry based methods developed so far have either used MRM based approaches in positive and negative mode or high resolution mass spectrometry based on accurate masses to characterise oxidised PC species(Hui et al., 2010, Gruber et al., 2012, Nakanishi et al., 2009, Davis et al., 2008). Furthermore, the primary oxidation products that are hydroperoxide species, which are very unstable and they get oxidatively fragmented to short chain oxidation products or are reduced to hydroxides by several enzymes in-vivo. Therefore, their abundance level in biological samples can be very low and can be below the detection level of existing methodologies. In our method, we tested a series of targeted approaches to selectively identify several oxPC species in a single run. The neutral loss scan for 34 Da in combination with precursor ion scanning for head group ions (selective for phospholipid class specific identification) allowed identification of several oxidised phospholipid species in a single run as demonstrated in our results. Moreover, the identification of mono-hydroperoxides, hydroxyl-hydroperoxides, and poly hydroperoxides was facilitated by using the targeted approach and in absence of specific standards.

Although, mass spectrometry technology provides wealth of information about your samples, it brings along the complexity of data analysis and intrinsic variability of instrumentation that can lead to misinterpretation of findings, unless the raw data is appropriately processed and normalised. Therefore, when the developed method is based on scanning approaches on a low resolution mass spectrometer, it is often required to validate the findings using a different chromatographic set-up or transferring the method to another high resolution mass spectrometry instrument. Therefore, we used an approach by transferring the methodology to a high resolution Q-TOF mass spectrometer to validate our findings. The method was also assessed for its qualitative and quantitative use by investigating the injection precision and inter-day variability. The relative standard deviation (CV %) was found to be less than 10 % for substantial number of species. There were some outliers that can be accounted for due to integration errors, peak distortion because of co-eluting species and extraneous level of oxidised species formed during storage.

The similar qualitative assessment was performed for the transferred method on the Q-TOF instrument to assess the technical variability and in-sequence variability. It showed comparable results to that observed on a low resolution Q-Trap instrument. Moreover, data analysis validation was performed by using different software (Progenesis QI) capable of automated pre-processing the raw mass spectrometry data to eliminate any non-biological variance, and also performs univariate and multivariate statistical analysis.

Once the robustness of the method was assessed, the method was employed for relative quantification of OxPC species in plasma samples of diabetic patients and healthy volunteers. We were able to identify several OxPC species in the plasma samples of diabetic and healthy patients that confirmed the sensitivity of our method. Moreover, the trend showed high abundance of OxPC species in diabetic patients when compared to healthy volunteers. However, when statistical analysis was performed for the data generated on high resolution mass spectrometer (Q-TOF), using progenesis QI software, the difference was found to be non-significant. This was on account of inherent biological variability found between abundances of OxPC species in plasma samples. This also may be explained by the complex model system such as plasma that was used for comparative analysis of OxPC species. Various biological factors may contribute to the levels of OxPC species in plasma samples and therefore, it raises the question of plasma being an appropriate model system for diabetic patients. Further investigation is required to match an appropriate model system to study diabetes as a disease for example: wound exudates of diabetic patients. Moreover, since our study was confined to only one class of phospholipid so far, it is possible that oxidised species of other classes of phospholipids may be up regulated. Type 2 diabetes being a metabolic disorder, it is very much possible that oxidised species of other phospholipids like cardiolipin and phosphatidylserine class, which are abundant in mitochondrial membrane may be formed preferentially due to oxidative stress.

Moreover, it is well established that glycated adducts with aminophospholipids like phosphatidylethanolamine and phosphatidylserines are upregulated in type 2 diabetes (Davies and Guo, 2014, Guo et al., 2010). These species cannot be identified using our method and this limits our analysis.

Overall, our work demonstrates an improved method using targeted approaches that can be used for comparative analysis of several OxPC species in biological samples.

## Supplementary Information

**Figure 1:**
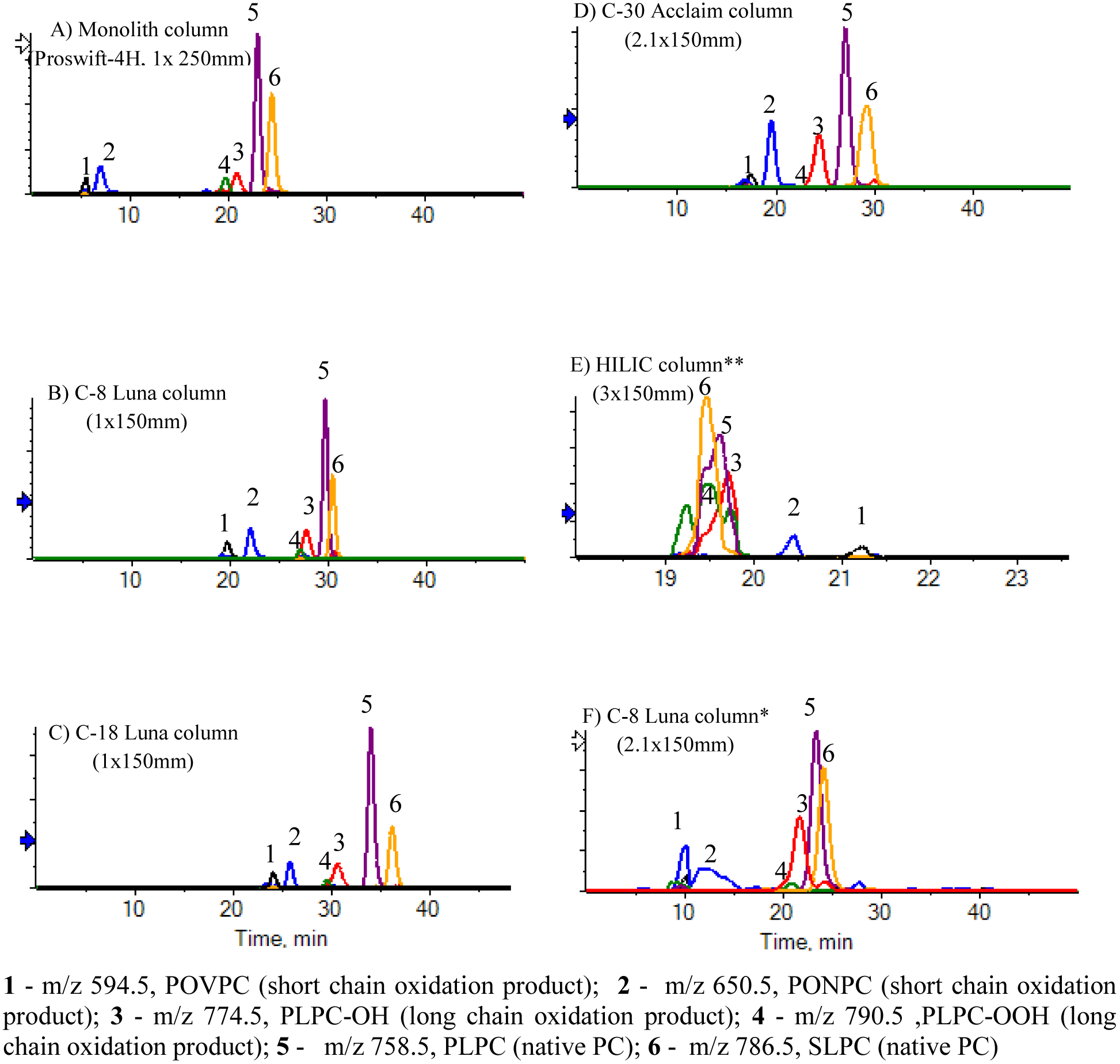
Extracted ion chromatogram (XIC) were generated of two representative species for each group; chain shortened products (POVPC,m/z 594.5; PONPC, m/z 650.5), long chain oxidation product (PLPC-OH, m/z 774.5; PLPC-OOH, m/z 790.5) and Native PCs (PLPC, m/z 758.5; SLPC, m/z 786.5) and overlaid to evaluate the elution profile of OxPCs using different column. Different solvent systems were also used to optimise the separation of different species. ** indicates solvent system for HILIC column: eluent A-20% Isopropyl alcohol (IPA) in Acetonitrile (ACN) and eluent B-20 % IPA in ammonium formate (20mM); * indicates another solvent system consisting of eluent A as Tetrahydrofuran (THF):methanol:water (30:20:50) and eluent B as THF: methanol: water (70:20:10) with 10mM ammonium acetate.

**Table 1:** Effect of flow rates on retention, sensitivity and resolution of representative oxidised species calculated by generating extracted ion chromatogram (XIC) and measuring peak area. The OxPC mixture was separated using monolith column at different flow rates and detection was performed on Q-trap mass spectrometer.

| Flow rate |  |  | 50 µl/minute |  | 100 µl/minute |  | 200 µl/minute |  |
| --- | --- | --- | --- | --- | --- | --- | --- | --- |
| compound | Monoisotopic mass (M+H)+ | Molecular Formula (neutral compound) | Retention time (in minutes) | Normalised Peak area (cps) | Retention time (in minutes) | Normalised Peak area (cps) | Retention time (in minutes) | Normalised Peak area (cps) |
| 16:0 lysoPC | 496.3402 | C24H50O7NP | 4.71 | 3.15E+06 | 2.25 | 2.82E+06 | 1.18 | 3.36E+05 |
| 18:0 lysoPC | 524.3710 | C26H54O7NP | 5.53 | 2.35E+08 | 2.66 | 1.29E+08 | 1.38 | 3.62E+07 |
| 18:2 lysoPC | 520.3407 | C26H50O7NP | 4.8 | 2.45E+05 | 2.34 | 3.53E+05 | 0 | 0.00E+00 |
| 18:1 lyso PC | 522.3561 | C26H52O7NP | 5.05 | 3.29E+07 | 2.44 | 2.13E+07 | 1.22 | 4.71E+06 |
| 20:4 lysoPC | 544.3403 | C28H50NO7P | 5.11 | 5.33E+05 | 0 | 0.00E+00 |  | 0.00E+00 |
| 13:0PC (internal standard) | 650.4755 | C34H68O8NP | 17.79 | 1.69E+07 | 10.94 | 1.07E+07 | 7 | 2.24E+06 |
| PLPC | 758.5694 | C42H80NO8P | 22.99 | 7.10E+08 | 15.83 | 6.72E+08 | 11.85 | 2.31E+08 |
| POPC | 760.5850 | C42H82NO8P | 23.52 | 1.61E+09 | 16.33 | 1.81E+09 | 12.32 | 6.73E+08 |
| SLPC | 786.6007 | C44H84O8NP | 24.45 | 5.49E+08 | 17.22 | 6.42E+08 | 13.21 | 1.44E+08 |
| SAPC | 810.6007 | C46H84NO8P | 24.77 | 5.86E+07 | 17.56 | 8.19E+07 | 13.54 | 2.18E+07 |
| P-7KPC(1palmitoyl-2-(7-oxoheptanoic acid) | 622.4079 | C31H60O9NP | 6.02 | 1.68E+07 | 2.89 | 1.12E+07 | 1.54 | 2.23E+06 |
| P-8KPC(1palmitoyl-2-(8-oxo-octanoic acid) | 636.4235 | C32H62O9NP | 6.4 | 4.01E+07 | 3.14 | 3.41E+07 | 1.6 | 7.26E+06 |
| P-7K5EPC (1palmitoyl-2-(7-oxo-5-heptenoic acid) | 620.3922 | C31H58O9NP | 6.28 | 4.48E+06 | 3.08 | 3.30E+06 | 1.53 | 6.08E+05 |
| P-11K9EPC(1-palmitoyl-2-(11-oxo-9-undecenoic acid)PC | 676.4548 | C35H66O9NP | 8.11 | 1.51E+07 | 4.08 | 1.02E+07 | 2.29 | 4.29E+06 |
| POVPC | 594.3763 | C29H56O9NP | 5.55 | 3.48E+06 | 2.66 | 2.55E+06 | 1.32 | 3.95E+05 |
| PONPC | 650.4391 | C33H64NPO9 | 6.96 | 1.48E+08 | 3.42 | 1.21E+08 | 1.79 | 2.33E+07 |
| S-7KPC | 650.4392 | C33H64O9NP | 9.69 | 2.91E+06 | 0 | 0.00E+00 | 0 |  |
| SOVPC | 622.4079 | C31H60O9NP | 7.55 | 2.22E+07 | 3.76 | 1.53E+07 | 2.02 | 3.10E+06 |
| SONPC | 678.4705 | C35H68NPO9 | 10.77 | 4.51E+07 | 5.57 | 2.99E+07 | 3.23 | 7.76E+06 |
| PAzPC | 666.4340 | C33H64O10NP | 6.86 | 5.37E+07 | 3.39 | 5.23E+07 | 2.38 | 5.12E+06 |
| SGPC | 638.4028 | C31H60O10NP | 7.48 | 1.47E+07 | 3.06 | 3.33E+07 | 1.56 | 6.43E+06 |
| SAzPC | 694.4653 | C35H68O10NP | 10.44 | 2.24E+07 | 5.45 | 1.53E+07 | 3.18 | 3.24E+06 |
| PLPC-OOH | 790.5593 | C42H80O10NP | 19.71 | 8.53E+07 | 12.99 | 7.02E+07 | 9.42 | 1.61E+07 |
| SLPC-OOH | 818.5911 | C44H84NO10P | 21.42 | 2.96E+07 | 14.54 | 3.87E+07 | 10.79 | 9.74E+06 |
| PEIPC | 828.5381 | C44H78O11NP | 18.55 | 2.88E+06 | 11.74 | 1.41E+06 | 5.17 | 1.58E+06 |
| PLPC(OH)-OOH | 806.5541 | C42H80NPO11 | 17.93 | 2.05E+07 | 11.21 | 1.31E+07 | 7.4 | 2.67E+06 |
| SLPC(OH)-OOH | 834.5855 | C44H84NO11P | 19.97 | 5.05E+06 | 13.24 | 4.60E+06 | 9.62 | 1.40E+06 |
| POPC-OH | 776.5800 | C42H82NO9P | 21.47 | 8.79E+07 | 14.5 | 8.90E+07 | 10.72 | 2.90E+07 |
| PLPC-OH | 774.5643 | C42H80O9NP | 20.86 | 1.29E+08 | 13.96 | 1.35E+08 | 10.25 | 4.19E+07 |
| SAPC-OH | 826.5956 | C46H84NO9P | 21.76 | 3.78E+06 | 12.42 | 2.53E+06 | 8.75 | 8.73E+05 |

**Table 2:**
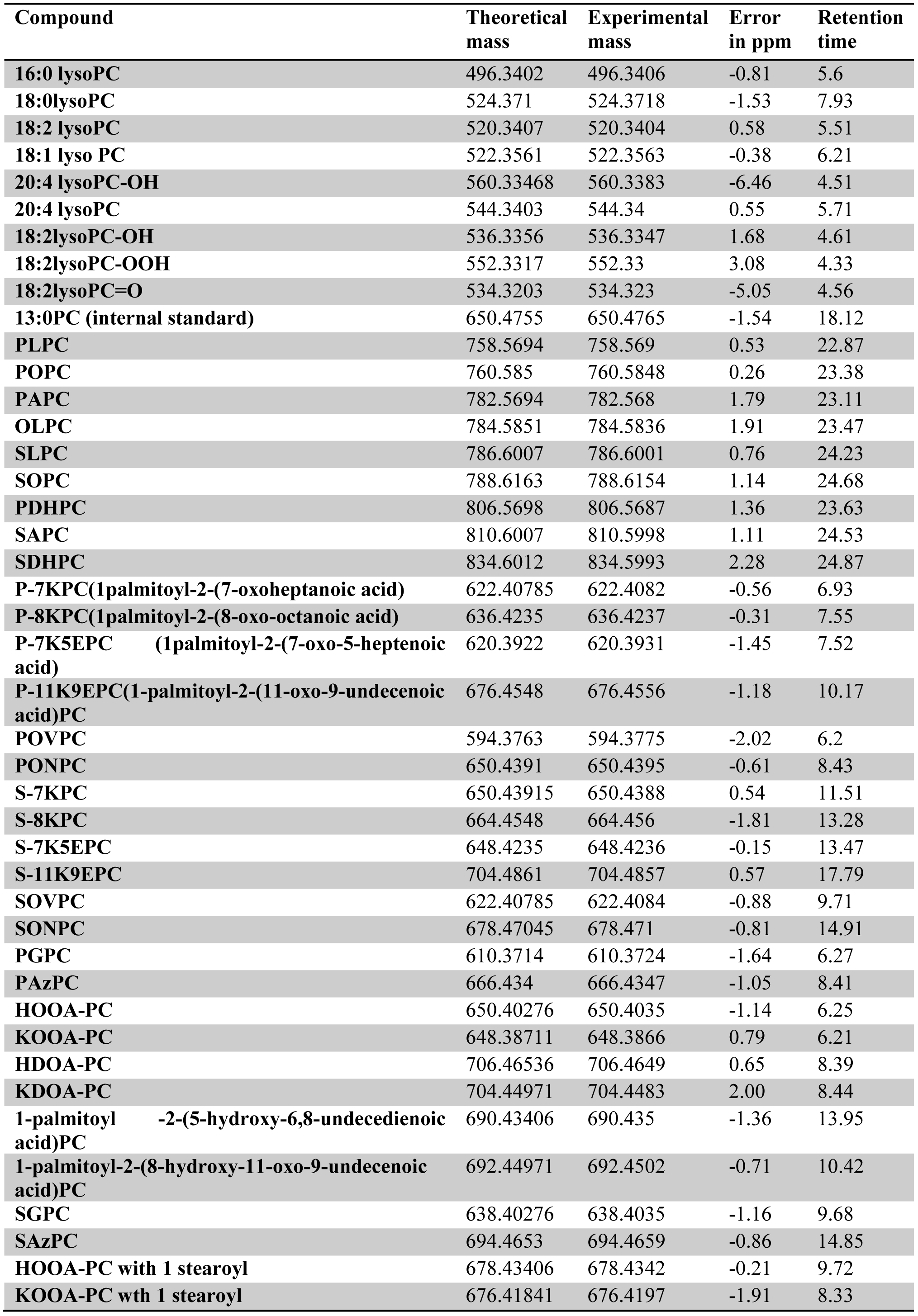
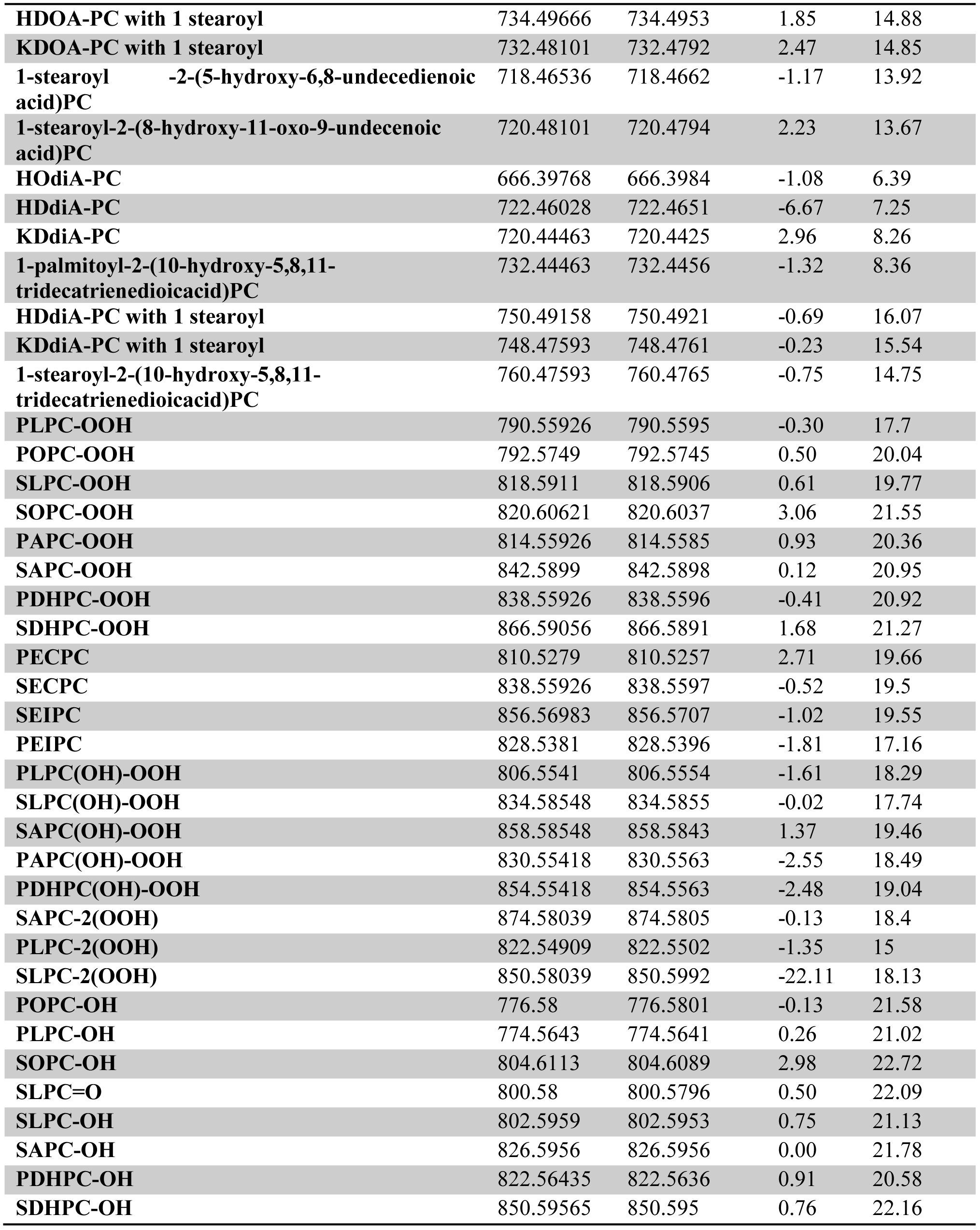
List of oxidised phosphatidylcholine (OxPC) species with accurate masses, experimental masses, measured retention time and molecular formula.

| Compound | Theoretical mass | Experimental mass | Error in ppm | Retention time |
| --- | --- | --- | --- | --- |
| 16:0 lysoPC | 496.3402 | 496.3406 | -0.81 | 5.6 |
| 18:0 lysoPC | 524.371 | 524.3718 | -1.53 | 7.93 |
| 18:2 lysoPC | 520.3407 | 520.3404 | 0.58 | 5.51 |
| 18:1 lyso PC | 522.3561 | 522.3563 | -0.38 | 6.21 |
| 20:4 lysoPC-OH | 560.33468 | 560.3383 | -6.46 | 4.51 |
| 20:4 lysoPC | 544.3403 | 544.34 | 0.55 | 5.71 |
| 18:2 lysoPC-OH | 536.3356 | 536.3347 | 1.68 | 4.61 |
| 18:2 lysoPC-OOH | 552.3317 | 552.33 | 3.08 | 4.33 |
| 18:2 lysoPC=O | 534.3203 | 534.323 | -5.05 | 4.56 |
| 13:0PC (internal standard) | 650.4755 | 650.4765 | -1.54 | 18.12 |
| PLPC | 758.5694 | 758.569 | 0.53 | 22.87 |
| POPC | 760.585 | 760.5848 | 0.26 | 23.38 |
| PAPC | 782.5694 | 782.568 | 1.79 | 23.11 |
| OLPC | 784.5851 | 784.5836 | 1.91 | 23.47 |
| SLPC | 786.6007 | 786.6001 | 0.76 | 24.23 |
| SOPC | 788.6163 | 788.6154 | 1.14 | 24.68 |
| PDHPC | 806.5698 | 806.5687 | 1.36 | 23.63 |
| SAPC | 810.6007 | 810.5998 | 1.11 | 24.53 |
| SDHPC | 834.6012 | 834.5993 | 2.28 | 24.87 |
| P-7KPC(1palmitoyl-2-(7-oxoheptanoic acid) | 622.40785 | 622.4082 | -0.56 | 6.93 |
| P-8KPC(1palmitoyl-2-(8-oxo-octanoic acid) | 636.4235 | 636.4237 | -0.31 | 7.55 |
| P-7K5EPC (1palmitoyl-2-(7-oxo-5-heptenoic acid) | 620.3922 | 620.3931 | -1.45 | 7.52 |
| P-11K9EPC(1-palmitoyl-2-(11-oxo-9-undecenoic acid)PC | 676.4548 | 676.4556 | -1.18 | 10.17 |
| POVPC | 594.3763 | 594.3775 | -2.02 | 6.2 |
| PONPC | 650.4391 | 650.4395 | -0.61 | 8.43 |
| S-7KPC | 650.43915 | 650.4388 | 0.54 | 11.51 |
| S-8KPC | 664.4548 | 664.456 | -1.81 | 13.28 |
| S-7K5EPC | 648.4235 | 648.4236 | -0.15 | 13.47 |
| S-11K9EPC | 704.4861 | 704.4857 | 0.57 | 17.79 |
| SOVPC | 622.40785 | 622.4084 | -0.88 | 9.71 |
| SONPC | 678.47045 | 678.471 | -0.81 | 14.91 |
| PGPC | 610.3714 | 610.3724 | -1.64 | 6.27 |
| PAzPC | 666.434 | 666.4347 | -1.05 | 8.41 |
| HOOA-PC | 650.40276 | 650.4035 | -1.14 | 6.25 |
| KOOA-PC | 648.38711 | 648.3866 | 0.79 | 6.21 |
| HDOA-PC | 706.46536 | 706.4649 | 0.65 | 8.39 |
| KDOA-PC | 704.44971 | 704.4483 | 2.00 | 8.44 |
| 1-palmitoyl -2-(5-hydroxy-6,8-undecdienoic acid)PC | 690.43406 | 690.435 | -1.36 | 13.95 |
| 1-palmitoyl-2-(8-hydroxy-11-oxo-9-undecenoic acid)PC | 692.44971 | 692.4502 | -0.71 | 10.42 |
| SGPC | 638.40276 | 638.4035 | -1.16 | 9.68 |
| SAzPC | 694.4653 | 694.4659 | -0.86 | 14.85 |
| HOOA-PC with 1 stearoyl | 678.43406 | 678.4342 | -0.21 | 9.72 |
| KOOA-PC with 1 stearoyl | 676.41841 | 676.4197 | -1.91 | 8.33 |
| **HDOA-PC with 1 stearoyl** | 734.49666 | 734.4953 | 1.85 | 14.88 |
| **KDOA-PC with 1 stearoyl** | 732.48101 | 732.4792 | 2.47 | 14.85 |
| **1-stearoyl -2-(5-hydroxy-6,8-undecdienoic acid)PC** | 718.46536 | 718.4662 | -1.17 | 13.92 |
| **1-stearoyl-2-(8-hydroxy-11-oxo-9-undecenoic acid)PC** | 720.48101 | 720.4794 | 2.23 | 13.67 |
| **HODiA-PC** | 666.39768 | 666.3984 | -1.08 | 6.39 |
| **HDdiA-PC** | 722.46028 | 722.4651 | -6.67 | 7.25 |
| **KDdiA-PC** | 720.44463 | 720.4425 | 2.96 | 8.26 |
| **1-palmitoyl-2-(10-hydroxy-5,8,11-tridecatrienedioic acid)PC** | 732.44463 | 732.4456 | -1.32 | 8.36 |
| **HDdiA-PC with 1 stearoyl** | 750.49158 | 750.4921 | -0.69 | 16.07 |
| **KDdiA-PC with 1 stearoyl** | 748.47593 | 748.4761 | -0.23 | 15.54 |
| **1-stearoyl-2-(10-hydroxy-5,8,11-tridecatrienedioic acid)PC** | 760.47593 | 760.4765 | -0.75 | 14.75 |
| **PLPC-OOH** | 790.55926 | 790.5595 | -0.30 | 17.7 |
| **POPC-OOH** | 792.5749 | 792.5745 | 0.50 | 20.04 |
| **SLPC-OOH** | 818.5911 | 818.5906 | 0.61 | 19.77 |
| **SOPC-OOH** | 820.60621 | 820.6037 | 3.06 | 21.55 |
| **PAPC-OOH** | 814.55926 | 814.5585 | 0.93 | 20.36 |
| **SAPC-OOH** | 842.5899 | 842.5898 | 0.12 | 20.95 |
| **PDHPC-OOH** | 838.55926 | 838.5596 | -0.41 | 20.92 |
| **SDHPC-OOH** | 866.59056 | 866.5891 | 1.68 | 21.27 |
| **PECPC** | 810.5279 | 810.5257 | 2.71 | 19.66 |
| **SECPC** | 838.55926 | 838.5597 | -0.52 | 19.5 |
| **SEIPC** | 856.56983 | 856.5707 | -1.02 | 19.55 |
| **PEIPC** | 828.5381 | 828.5396 | -1.81 | 17.16 |
| **PLPC(OH)-OOH** | 806.5541 | 806.5554 | -1.61 | 18.29 |
| **SLPC(OH)-OOH** | 834.58548 | 834.5855 | -0.02 | 17.74 |
| **SAPC(OH)-OOH** | 858.58548 | 858.5843 | 1.37 | 19.46 |
| **PAPC(OH)-OOH** | 830.55418 | 830.5563 | -2.55 | 18.49 |
| **PDHPC(OH)-OOH** | 854.55418 | 854.5563 | -2.48 | 19.04 |
| **SAPC-2(OOH)** | 874.58039 | 874.5805 | -0.13 | 18.4 |
| **PLPC-2(OOH)** | 822.54909 | 822.5502 | -1.35 | 15 |
| **SLPC-2(OOH)** | 850.58039 | 850.5992 | -22.11 | 18.13 |
| **POPC-OH** | 776.58 | 776.5801 | -0.13 | 21.58 |
| **PLPC-OH** | 774.5643 | 774.5641 | 0.26 | 21.02 |
| **SOPC-OH** | 804.6113 | 804.6089 | 2.98 | 22.72 |
| **SLPC=O** | 800.58 | 800.5796 | 0.50 | 22.09 |
| **SLPC-OH** | 802.5959 | 802.5953 | 0.75 | 21.13 |
| **SAPC-OH** | 826.5956 | 826.5956 | 0.00 | 21.78 |
| **PDHPC-OH** | 822.56435 | 822.5636 | 0.91 | 20.58 |
| **SDHPC-OH** | 850.59565 | 850.595 | 0.76 | 22.16 |

